# Profiling miRNAs involved in Human Oligodendrocyte Precursor Cell Differentiation and Maturation

**DOI:** 10.64898/2025.12.25.696416

**Authors:** Mansoureh Barzegar, Asmita Dhukhwa, Vaidehi Nilesh Patel, Fernanda C Velasquez, Samarjit Das, Arun H Patil, Marc K. Halushka, Donald J. Zack, Xitiz Chamling

## Abstract

MicroRNAs (miRNAs) are evolutionarily conserved post-transcriptional regulators that play critical roles in cellular development and differentiation across species. Although the importance of miRNAs in oligodendrocyte lineage cell (OLLC) differentiation has been extensively studied in rodent models, their roles in human OL development remain less understood. To address this gap, we used a human embryonic stem cell (hESC) reporter system designed to study human OLs and OL progenitor cells (OPCs). Using an optimized differentiation protocol, we used the reporter hESCs to generate and isolate well-characterized OLLCs at specific developmental stages and performed next-generation sequencing-based miRNA profiling to identify stage-specific miRNAs enriched during OL lineage specification and maturation. In addition to canonical miRNAs known to be enriched at various stages of OL development, our study identified several lesser-known miRNAs with distinct stage-specific enrichment patterns that may serve as useful molecular markers for classifying human CNS cell types in future studies. Target analysis of OPC-and OL-enriched miRNAs revealed key genes, including transcription factors ZNF488 and DLX1, cytoskeletal regulator CSNK2B, and potassium channel gene KCNJ1, along with key signaling pathways such as AKT, SMAD2/3, estrogen receptor, and insulin signaling, which regulate OPC and OL lineage function. These findings advance our understanding of the OLLC-specific miRNAs, and miRNA-mediated regulatory networks governing human OL differentiation and maturation and provide promising therapeutic targets for future studies aimed at restoring myelin integrity and improving outcomes in demyelinating diseases.

## Introduction

Oligodendrocytes (OLs) are the specialized glial cells in the central nervous system (CNS) responsible for forming the myelin sheath that wraps around neuronal axons. The myelin is crucial for the efficient transmission of electrical impulses along neurons. During development, neural progenitor cells (NPCs) in the CNS differentiate into oligodendrocyte progenitor cells (OPCs), which migrate throughout the brain and spinal cord. The OPCs then terminally differentiate into immature OLs. Upon receiving appropriate environmental cues, these immature OLs further mature, wrapping around nearby receptive axons and myelinating them^1^. Myelination of neuronal exons is essential for saltatory conduction of action potential, providing trophic support to the axons, and maintaining their survival and function. Consequently, loss of myelin integrity could lead to severe neurological disorders, such as Multiple Sclerosis, inherited Leukodystrophies, and optic neuritis-related to vision loss. To facilitate the repair of damaged myelin and maintain myelin integrity, it is important to understand the molecular mechanisms that regulate OL differentiation and maturation. The process of fate specification at NPC and OPC stages, as well as OL development and maturation, are controlled by several extrinsic and intrinsic factors, which regulate changes at both transcriptional as well as post transcriptional levels. MicroRNAs (miRNAs) have been recognized as one of the key post-transcriptional regulators involved in determining the fate of NPCs and their transition into OPC and OL lineage^2,3^.

MicroRNAs (miRNAs) are small, noncoding RNA molecules that are trimmed from longer hairpin-loop RNA sequences into functional 19-21 mers by the miRNA processing enzymes DICER1 and DROSHA^4^. miRNAs recognize a complementary sequence in the 3′ UTR of a protein-coding messenger RNA (mRNA) and inhibit expression of specific proteins, either by repressing RNA translation or directly promoting the degradation of the associated mRNAs^5^. Each miRNA can target multiple mRNAs, allowing them to modulate numerous pathways, and by regulating molecular pathways, miRNAs have the potential to modulate cell proliferation, differentiation, and lineage specification^6^.

miRNAs play a critical role in the development of several CNS cell types, including neurons, astroglia, and oligodendroglia, and aberrant expression of miRNAs have been implicated in a broad range of demyelinating and neurodegenerative diseases^7^. The importance of miRNA in OL development and myelination has been well documented, especially by a study where deletion of Dicer in OPCs led to delayed OL differentiation and myelination^8,9^. This and several other studies have identified and confirmed the significance of specific miRNAs, such as miR219 and miR338, in OPC and OL differentiation and maturation^9^. However, most miRNA profiling and biochemical studies aimed at understanding the role of miRNAs in OPC and OL development have been performed using murine models. Although human and mouse OPCs and OLs are similar, there are important differences^10-12^, and detailed profiling and characterization of miRNAs that regulate the differentiation of human OL is still lacking.

One of the main challenges in using human OPCs and OLs (hOPCs/OLs) for such studies is the limited access to primary human cells, especially for studies requiring cells at different developmental stages. While hOPCs/OLs differentiated from human pluripotent stem cells (hPSCs) offer a promising alternative to address this limitation, obtaining sufficient numbers of well-characterized human hPSC-derived OL lineage cells (OLLCs) remain challenging. Due in part to this constraint, to our knowledge, only one study profiling global miRNAs in human hESC-derived OLLCs has been reported to date^13^. Although that study identified several interesting miRNAs, it was limited by the available miRNA profiling technology and the quality of OPC differentiation protocol at the time^13^.

In this study, we used our robust method to generate well-characterized OPCs and capture miRNAs at seven different stages of OLLC differentiation. We utilized our CRISPR-engineered hESC line, which enables scalable differentiation and enrichment of human OPCs at different developmental stages^14,15^. The reporter hESC was differentiated into OL lineage and then NPCs, OPCs, and OLs were selectively isolated at different stages of differentiation. We then performed next-generation sequencing (NGS)-based miRNA profiling, which not only identified miRNAs previously shown to be important for mouse OPC differentiation, but also revealed a number of additional miRNAs that are distinctly enriched in differentiating human OPC and OL populations.

## Results

### miRNA isolation from hESC derived cells at different stages of oligodendrocyte differentiation

To comprehensively profile miRNA expression at various stages of human OLLC development, we utilized a hESC (RUES1) reporter system that we developed in our laboratory^14,15^ (Figure1A). As previously reported^15^, this reporter cell line, referred to as PPM (PDGFRα-PLP1-MBP reporter) hereafter, consists of three reporters: **1)** PDGFRα-P2A-tdTomato-P2A-Thy1.2, where endogenous *PDGFRα* expression leads to the production of tdTomato fluorescent marker and cell-surface protein, Thy1.2. The tdTomato protein localizes to the cytoplasm whereas Thy1.2 localizes to the cell surface, enabling the immunopurification of PDGFR*α*-expressing cells using anti-Thy1.2 antibody; **2)** PLP1-sfGFP, in which expression of sfGFP is driven by an early OL marker *PLP1*; and **3)** MBP-P2A-secNLuc, where NLuc protein is secreted into the cell culture media, enabling serial sampling of the media for quantitative analysis of MBP expression^15^.

We differentiated three hESC clonal lines of this reporter system into hOPC/OL following our established protocol^14,15,16^ and collected cell populations at defined stages: stem cells (SC) at day 0, neural stem cells (NES) at day 8, neural progenitor cells (NPC) at day 20, and early OPCs at day 40 (Figure 1B, C). At day 75, post mitotic PDGFRα-tdTomato+ OPCs were purified using magnetic-activated cell sorting (MACS) with Thy1.2 antibody conjugated microbeads. Using this MACS method, we regularly achieve >95% tdTomato+ OPC population as confirmed with flow cytometry analysis (Figure 1D). On day 95 of differentiation, oligodendrocytes were enriched using O4 microbeads, which recognizes membrane sulfatide highly specific to early stage OLs. Although it is possible to sort OLs based on PLP1-GFP expression using fluorescence activated cell sorting (FACS), we have found that FACS is quite harsh on OLs, resulting in significant cell death and poor RNA quality from sorted cells. Therefore, we prefer to use MACS-based cell purification whenever possible. Next, we verified the identity of our different cell populations by assessing the expression of cell-type-specific markers using qPCR analysis: *OCT4* for SC, NES and *SOX2* for NPCs, *OLIG2* and *PDGFRα* for early OPCs, and *MBP* and *PLP1* for OLs (Figure 1E).

**Figure 1.**
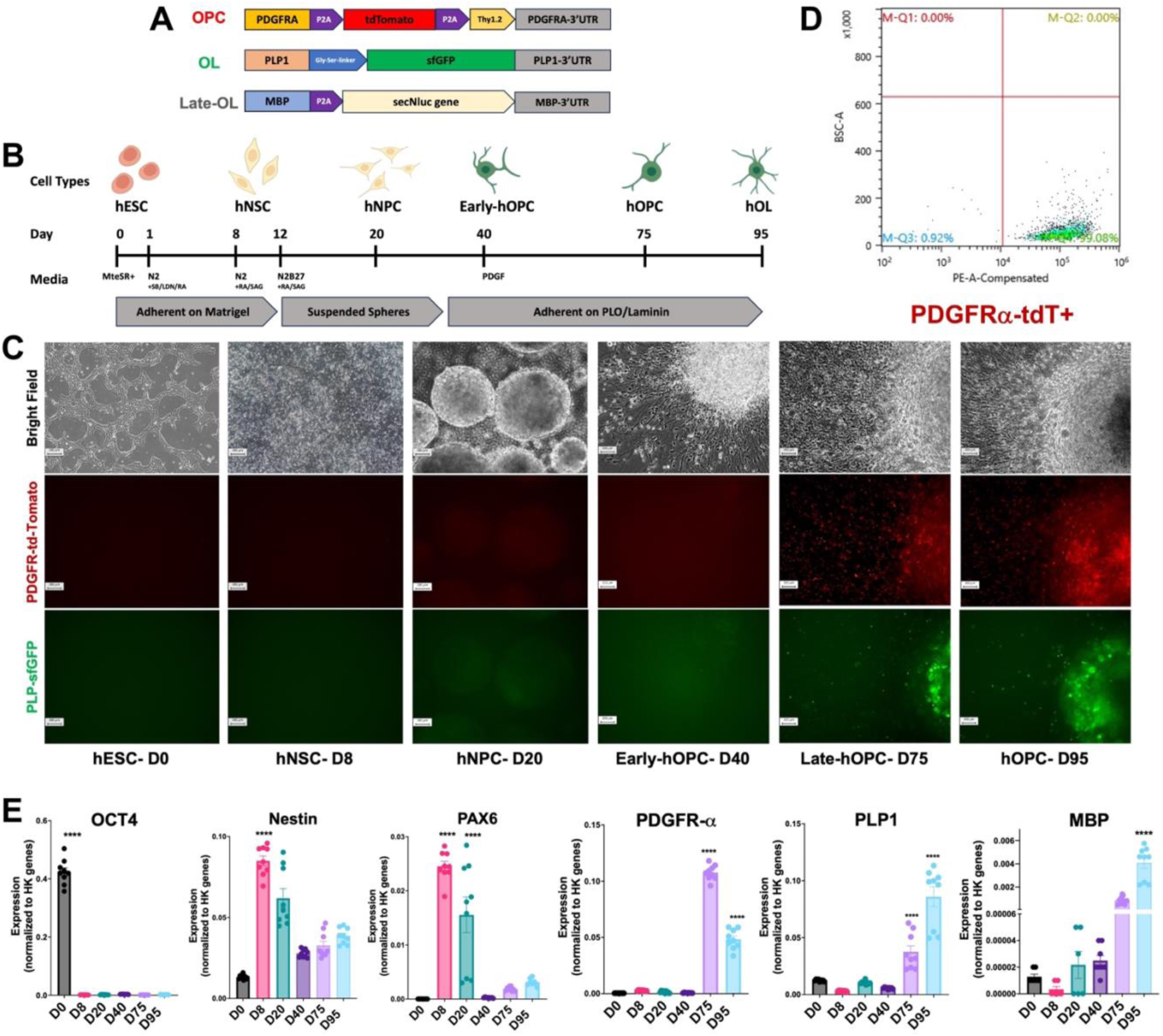
miRNA isolation from a reporter hESC derived cells at different stages of oligodendrocyte differentiation. **A)** Schematic of the CRISPR-Cas9-engineered hESC reporter cell line (PPM). **B)** Timeline showing differentiation and maturation of hOPCs and hOLs derived from PPM cell line. **C)** Representative images of stage-specific cell population during the differentiation process. **D)** An example of flow cytometry analysis showing 99% tdTomato+ hOPCs following purification using Thy1.2 antibody conjugated magnetic microbeads. **E)** qPCR confirming the expression of stage-specific markers in each cell population. Three clonal lines were used as biological replicates. Statistical significance was calculated using one-way ANOVA, *p<0.05, **p<0.01, ***p<0.001, ****p<0.0001, ns>0.05. Data represents mean values ± S.E.M.

### Next generation sequencing-based miRNA profiling

miRNA profiling has traditionally been performed using microarray platforms, NanoString nCounter system, or quantitative real-time PCR, each of which has limitations that impact their ability to provide a broad and accurate quantification of miRNAs^17^. Recent advancements in next-generation sequencing (NGS) methods offer a more advantageous approach to miRNA profiling compared to these other platforms^18^, as the NGS-based method is not limited by predefined features, probe design, or array background^19^. NGS is not only suitable for confirming the known miRNAs, similar to qRT-PCR and microarray platforms, but it can also detect novel miRNAs. Therefore, we opted to use the Qiagen miRNA library prep and Illumina Hiseq for this miRNA profiling study (Figure S1).

This NGS-based approach efficiently captured mature miRNA, with approximately 50% of the reads (∼40 million of the ∼80 million trimmed reads from the 20 runs) mapping to mature human miRNA in miRbase (Figure 2A, Table S1 and S2). An average of 657 mature miRNAs were identified across each cell population, with the median range between 391 and 1017 miRNAs (Figure 2B). The highest number of miRNAs were captured in the D20 NPC population, while the O4+ OLs had the least (Figure 2B). Principle component analysis (PCA) showed clear separation of clusters corresponding with different time points, highlighting that temporal changes in miRNA expression are strongly associated with the differentiation process. When compared to the SC stage, the most pronounced separation was observed in more mature cells at day 75 and day 95 (D75 OPCs, D75 non-OPC (flow through; FT), and D95 O4+ OL). Among the more mature population, OPCs (day 75) and OLs (day 95) clusters were closer to each other than the non-OPC population (D75-FT) (Figure 2C). The heatmap of differentiation stage-specific differentially expressed miRNAs further underscores these findings, revealing approximately 900 miRNA that are differently expressed during the differentiation process (Figure 2D). The majority of miRNA that were highly expressed during SC stage showed a progressive reduction as the cell matured, and vice versa. This pattern suggests, as expected, that miRNA expression profiles change more dramatically during the differentiation process than between OPC/OL and non-OPC lineage cells at similar time-points. Our NGS-based miRNA profiling provides a comprehensive overview of miRNA dynamics during human OL lineage development, revealing key miRNAs that potentially regulate maturation and lineage specification at various stages.

**Figure 2.**
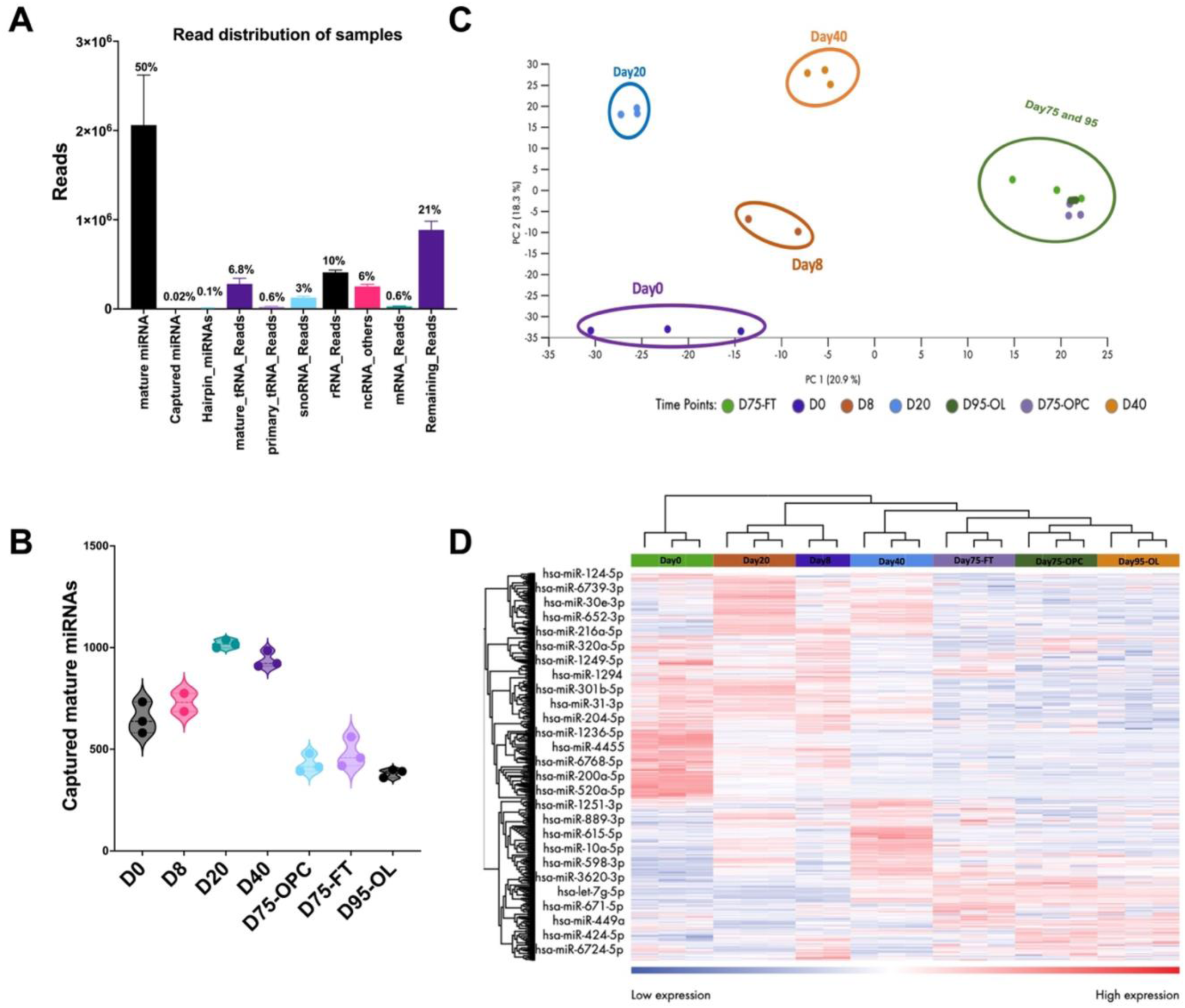
The NGS-based approach efficiently captured mature miRNA. **A**) Distribution of various small RNA species captured across samples. The Y-axis represents a number of sequencing reads, and X-axis indicates different types of small RNAs detected. The percentage of filtered reads corresponding to each small RNA type is displayed above each bar. **B)** Distribution of the number of mature miRNAs captured at different time points during the differentiation process. **C)** Principal Component Analysis (PCA) of miRNA expression profiles across distinct differentiation time points. Samples are represented as points, and their clustering reflects similarities or differences in miRNA expression patterns during OL differentiation. **D)** Heatmap of differentially expressed miRNAs during different stages of OL development. Rows represent individual miRNAs, and columns correspond to differentiation time points. Expression levels are visualized using a color scale, with statistical significance set at false discovery rate (FDR) < 0.01.

### Stage-specific enrichment of previously characterized miRNAs during OL development

As a validation of our miRNA data, we assessed the presence of miRNAs known to be upregulated in various cell types. As expected, miR-302b-3p, which is known to be upregulated in human pluripotent stem cells (hPSCs)^20,21^ and has been reported to promote reprograming of human somatic cells to hPSCs^22^, was highly enriched in the hESCs but not in other populations (Figure 3A). The expression of miR-26b-5p^23^ and miR-103a-3p^24^, which have been reported to initiate ESC to NPC differentiation and whose continued enrichment leads to neural differentiation in murine ESC, were enriched in our D20 NPCs and D40 progenitor cells, with its expression was reduced in the OPC and OL population (Figure 3B).

**Figure 3.**
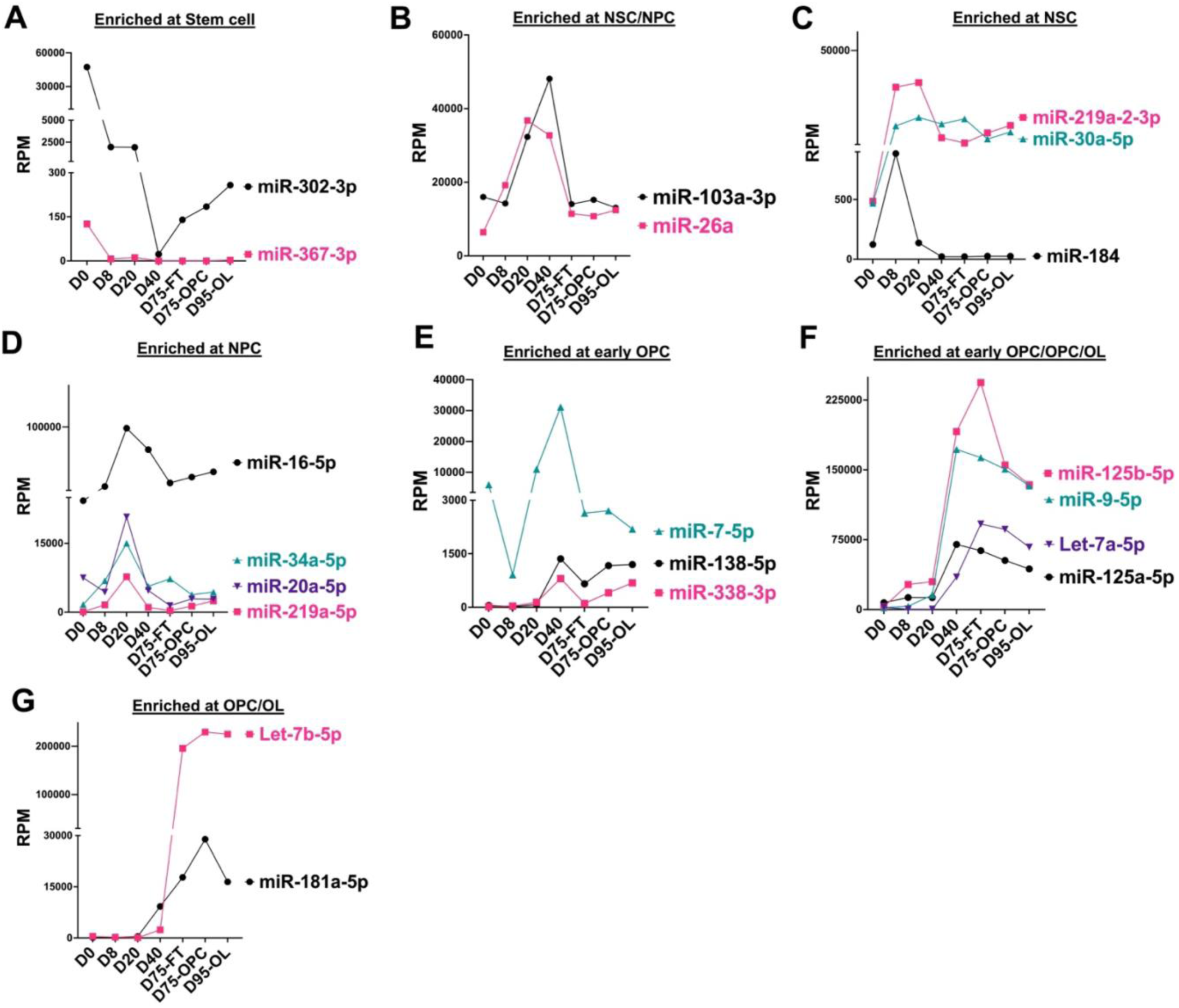
Stage-specific enrichment of previously characterized miRNAs during OL development. Each graph illustrates the expression patterns of miRNAs enriched at specific developmental stages: **A)** Day 0 representing the pluripotent stem cell stage. **B)** Neural stem cells (NSCs) and neural progenitor cells (NPCs) at Days 8 and 20. **C)** NSCs at Day 8. **D)** NPCs at Day 20. **E)** Early OPCs at Day 40. **F)** Early OPCs, mature OPCs, and OLs. **G)** Mature OPCs and OLs at Days 75 and 95. Each data point represents the average reads per million (RPM) from three biological replicates.

The miR-219 family members, namely miR-219a-2-3p and miR-219a-5p were enriched in OPCs and OLs compared to non-OPC population; however, their highest expression was detected in the D20 NPCs when compared to other time-points (Figures 3C-D). Similar to previous reports^25,26^, this data suggests that continued expression of miR-219a-5p and miR-219a-2-3p is crucial to NPC differentiation and important for OL lineage specification. Similarly, miR-138, and miR-338-3p, which have been implicated in OPC differentiation and OL maturation^8,25-27^ were enriched in D40 precursors, OPCs and OLs, with their expression significantly reduced in the non-OPC population (Figure 3E).

We also compiled a list of approximately 150 miRNAs previously reported to be involved in OPC and OL differentiation and maturation (Table S3) and assessed their expression in our dataset. Since most of these reports are based on rodent models, we observed only limited overlap with our data. However, stage-specific enrichment of several miRNAs was conserved between the previous reports and our dataset. Notably, enriched expression of miR-7-5p, miR-16-5p, miR-20a-5p, and miR-34a-5p in NPCs and early OPC was also observed in our dataset, suggesting their involvement in the initiation of OL differentiation^28,29,30^ (Figures 3D-E). Several other miRNAs, including miR-9-5p^31,32,33^, miR-125b-5p, miR-125a-5p^34^, and let-7a-5p, also showed increased expression in the hOPC (day 75 population) in our dataset (Figure 3F). Notably, let-7b-5p and miR-181a-5p^8,35,36^ showed significantly increased expression at OPC stage, which remained elevated at later time point (OL, day 95) (Figure 3G). These data further validate that our reporter hESC-derived cells and the miRNA profiling method can effectively capture mature miRNAs involved in OLLC development.

Nonetheless, it is worth noting that several miRNAs previously implicated in hESC differentiation to OL lineage^13^ did not show expression in the expected populations in our data. For example, miR-184, which is reported to be specific to OL^2,13^ was enriched only in the NPCs but significantly reduced in OPCs and OLs (Figure 3C); miR-367 that is reported to be enriched in hESC, NPCs and OPCs^20,37^ had a higher expression in hESCs compared to NPCs but it was not detected in the other populations (Figure 3A); and miR-663, miR-1225-5p, miR-638 that were previously reported to be enriched in hESC-derived OL^13^, were surprisingly not detected in any of the cell population in our dataset.

### Identification of differentiation stage-specific enriched miRNAs

To identify miRNAs with marked differential expression across different OLLC differentiation stages that have not been previously reported, we calculated the variance in miRNA expression across various time points and selected the 50 miRNAs with the highest variance (top 2nd percentile) across different time-points (Figure 4A). This analysis revealed several known miRNAs, as well as a few novel miRNAs, that are enriched in a stage-specific manner. For example, in addition to miR-302b-3p and miR-302a-3p, which were previously reported, we found a highly specific and significant enrichment of miR-183-5p and miR-182-5p at the stem cell (SC, day 0) stage (Figure 4B). miR-483-3p was specifically enriched at the neural stem cell (NSC, day 8) stage (Figure 4C), while upregulation of miR-454-3p, miR-363-3p, miR-151a-3p, and miR-10b-5p were highly specific to the neural progenitor cell (NPC, day 20) stage (Figure 4D). Additionally, miR-135a-5p, miR-27b-3p, miR-342-3p, miR-99b-5p, and miR-30d-5p exhibited specific enrichment during the pre-OPC stage (day 40) (Figure 4E). We also identified two miRNAs, miR-1226-3p, and miR-383-5p specifically enriched in OPC (D75) population (Figure 4F). A drastic increase in the expression of miR-9-3p was observed during differentiation, with OPC specific further enrichment (Figure 4F).

**Figure 4.**
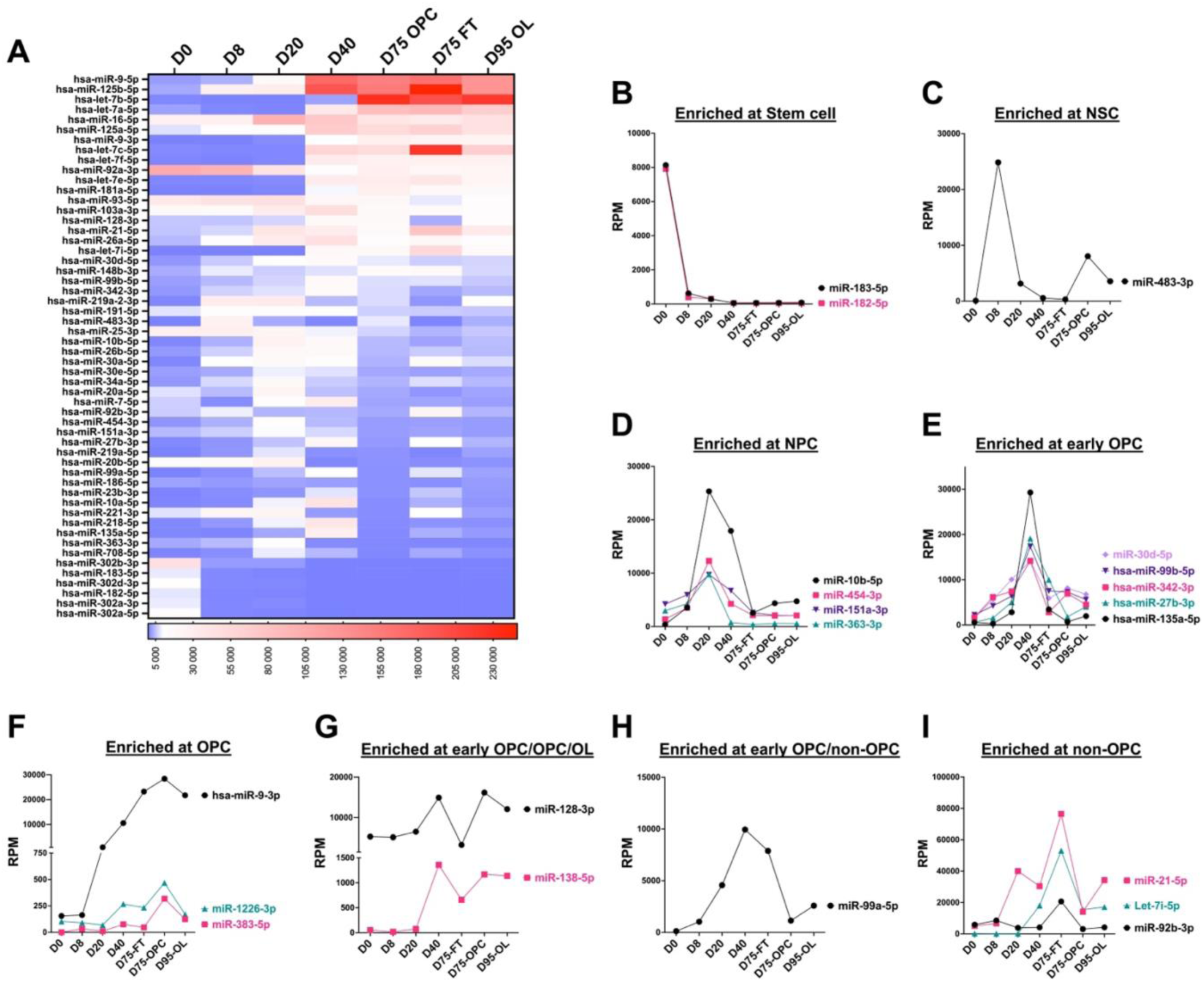
Variance analysis reveals stage-specific enrichment of miRNAs during OL differentiation. **A)** Heatmap showing the top-variance miRNAs across OPC to OL differentiation. **B–I)** Graphs illustrating the stage-specific enrichment of miRNAs uniquely identified in our dataset. **B)** Expression pattern of miRNAs enriched in pluripotent stem cell population (Day 0; **C)** NSC (Day 8); **D)** NPC (Day 20); **E)** early OPC (Day 40; **F)** OPC (Day 75); **G)** Early OPC/OPC/OL (Day 40, 75 and 95); **H)** early OPC and non-OPC (Day40 and flow through from day 75); and **I)** only non-OPC (flow through from day 75. Each data point represents the average normalized RPM from three biological replicates.

The variance analysis also highlighted two miRNAs, miR-138-5p and miR-128-3p, which are enriched in the early-OPC population (day 40), with this enrichment persisting in the OPC and OL populations (Figure 4G). When compared to the OPC/OL population, the expression of both miRNAs, particularly that of miR-128, was significantly reduced in the non-OPC populations, suggesting their importance in maturation of neural stem cells into OLLCs. On the other hand, miR-99a-5p was upregulated in early OPC and non-OPC (day 75 FT) populations (Figure 4H), while miR-92b-3p, let-7i-5p, and miR-21-5p were enriched only in the non-OPC population (Figures 4I). Since our previous study demonstrated that the flow-through after OPC immune-purification primarily comprises astrocyte cells^15^, these data suggest a potential role for these miRNAs in the specification of astrocytes from NPCs. Overall, this analysis helped generate a list of human OLLC differentiation stage specific marker miRNAs.

### miRNAs that are differentially expressed in human OPC and OLs population

Since OPC to OL differentiation is inhibited in MS and other demyelinating diseases, we were particularly interested in identifying miRNAs that are potentially involved in OPC formation and their maturation into myelinating OLs. Therefore, we performed differential expression analysis among three mature cell populations: OL vs OPC, OPC vs non-OPCs, and OLs vs non-OPC samples (Figures 5A-C). This analysis revealed 52 (26 up /26 down), 129 (40 up / 89 down), and 106 (42 up/ 64 down) differentially expressed miRNAs (FC >1.5 and FDR p-value<0.05) in the OL vs OPC, OPC vs non-OPC, and OLs vs non-OPC datasets respectively (Figures 5D, E). Interestingly, only miR-219-5p was consistently upregulated and only miR-1247-5p was consistently downregulated across all three datasets, highlighting the potential roles of miR-219-5p upregulation and miR-1247-5p downregulation in OPC lineage selection and OL maturation.

**Figure 5.**
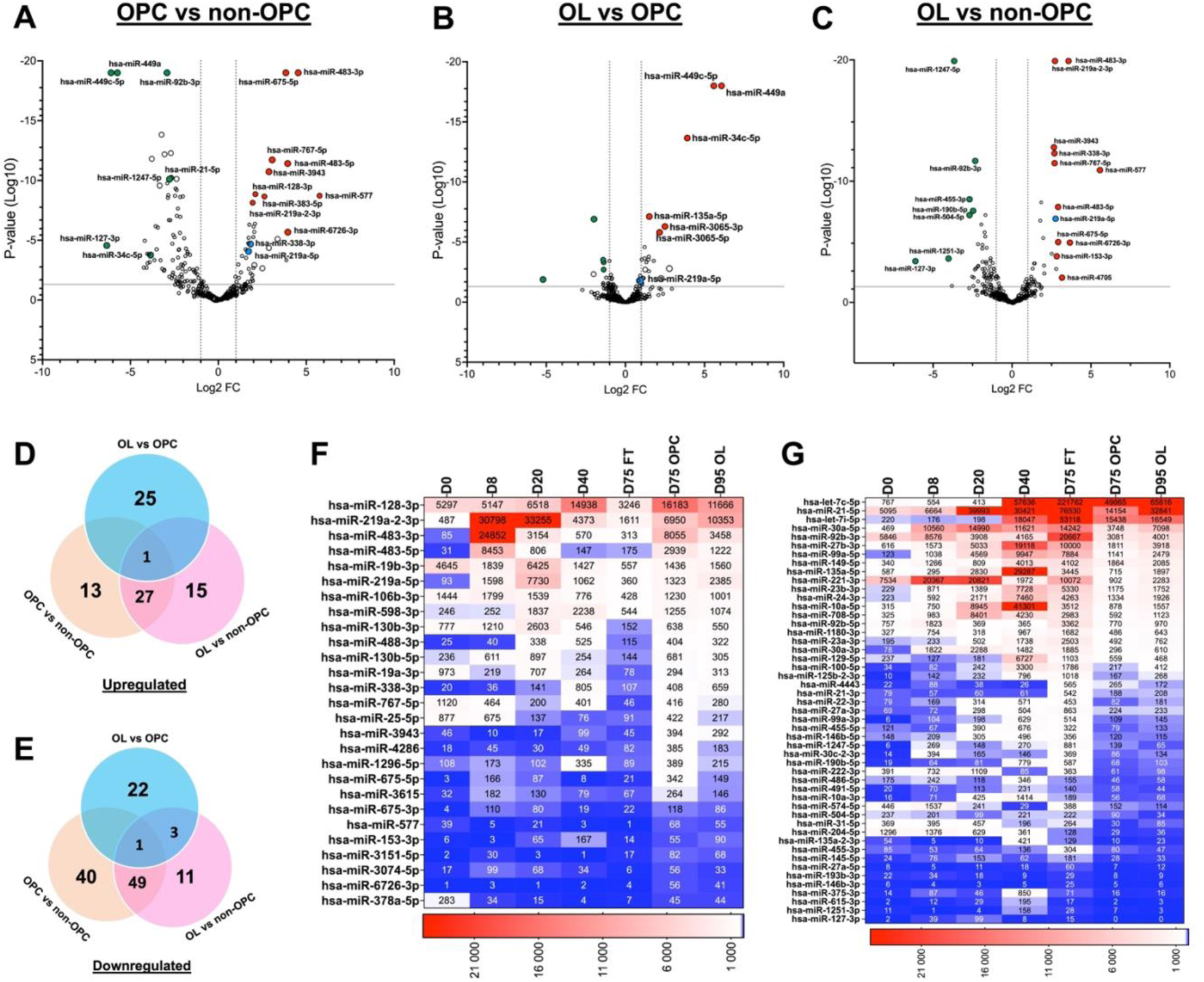
OPC- and OL-specific miRNA enrichment analysis. Volcano plot showing differentially expressed miRNAs between: **A)** OPC and non-OPC, **B)** OL and OPC, and **C)** OL and non-OPC populations. A log2 fold change > 1 or < -1 with P-value<0.01 is considered significant. Venn diagrams illustrating **D)** upregulated and **E)** downregulated miRNAs across the three comparisons. miR-219-5p is identified as the commonly upregulated miRNA, while miR-1247-5p is the commonly downregulated miRNA across all comparisons. Heatmap of **F)** 27 upregulated and **G)** 49 downregulated miRNAs from the comparisons of OPCs vs non-OPCs and OLs vs non-OPCs populations highlight miRNAs specifically enriched in OPC/OL populations.

Among the miRNAs upregulated in day 95-OL when compared to day 75-OPC are miR-449c-5p, miR-449a, and miR-34c-5p. Additional miRNAs such as miR-135a-5p, miR-3065-3p, miR-3065-5p, and miR-221-5p, were also upregulated in the OL when compared to OPC, although to a lesser extent (Figures 5B). Interestingly, many of those miRNAs enriched in the OL population, including miR-449c-5p, miR-449a, and miR-34c-5p, were also enriched in non-OPC (flow-through; FT) population when comparing OPCs vs non-OPCs (Figures 5A). This observed overlap in miRNA enrichment may be explained by theinadvertent inclusion of contaminating cells in a sample. For example, during thy1.2 MACS purification, which selectively isolates PDGFRα-tdTomato expressing OPCs, a small population of mature OLs may remain in the flow-through, leading to higher expression of OL associated miRNAs in non-OPC samples. Similar inadvertent contamination could occur in the D95 OL population, which is isolated using O4 antibodies. Although this method selects cells significantly enriched for OL markers (Figure 1E), it may also contain a small fraction of astrocytes, another major cell type in our differentiating culture system^15^. Since astrocytes constitute a significant proportion of the non-OPC FT sample, miRNAs commonly enriched in both OLs and non-OPC may instead represent astrocyte-enriched miRNAs.

Therefore, to identify miRNAs specifically associated with OPC/OL populations, we focused on candidates enriched in both OPCs and OLs when compared to non-OPC populations. In this analysis, 27 miRNAs were commonly upregulated, and 49 miRNAs downregulated in both OPC vs non-OPC and OL vs non-OPC datasets (Figures 5D-G). Among the miRNAs enriched in OPCs and OLs were miR-219a-5p and miR-338-3p, two widely studied miRNAs known for their roles in OL differentiation and myelination, further validating the robustness of our method and our data. Similarly, miR-128 expression, which was initially upregulated in day 40 pre-OPCs, remained enriched in OPCs and OLs but not in non-OPC population (Figure 5F). In line with our findings, miR-128-3p was recently reported to reverse fibrinogen-mediated inhibition of OPC differentiation and remyelination following cerebral ischemia^38^.

In addition, we identified several miRNAs that have not been reported or extensively studied in OPC/OL biology. Notably, at least eight miRNAs (miR-3943, miR-1296-5p, miR-4286, miR-3615, miR-675-5p, miR-3151-5p, 6726-3p, and miR-577) were specifically enriched during the OPC and OL stages, suggesting their important role in OLLC differentiation (Figure 5F). Conversely, among the downregulated miRNAs, miR-92b-5p/3p, miR23a-3p, miR4443, and miR-21-3p were among the candidates specifically enriched in the non-OPC FT population (Figure 5G). The stage-specific enrichment of these miRNAs suggests their involvement in OPCs lineage maintenance and their eventual differentiation into oligodendrocytes. These findings also emphasize the dynamic regulation of miRNAs during differentiation and highlight stage-specific candidates that are likely to play crucial roles in OLLC fate determination.

### Biological pathways and molecular networks regulated by differentially expressed miRNAs

To investigate potential biological pathways and molecular networks regulated by differentially expressed miRNAs, we performed Ingenuity Pathway Analysis (IPA) for each comparison groups: OPC vs non-OPC, OL vs non-OPC, and OL vs OPC (Figures 6A-B and S2). A substantial proportion of OPC/OL enriched miRNAs, as well as several miRNAs downregulated in comparison to non-OPCs (FT) were associated with predicted activation of AKT, PI3K, and Insulin signaling pathway, and with the inhibition of SMAD2/3 pathway. Additionally, inhibition of the estrogen receptor pathway was observed in the OL vs non-OPC comparison (Figure 6B), which aligns with previous studies reporting that pharmacological inhibition of this pathway enhances OL maturation.

**Figure 6.**
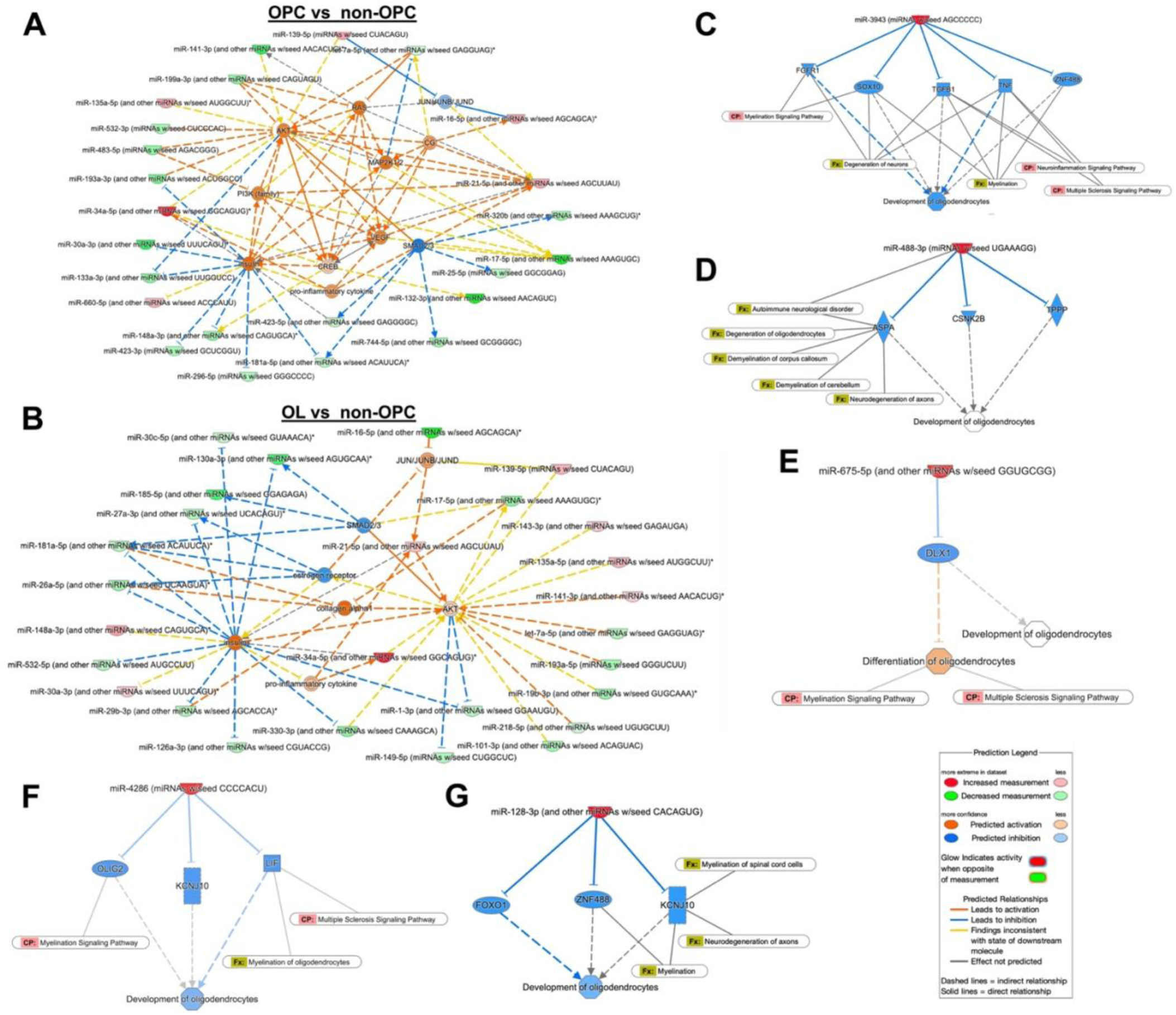
Ingenuity Pathway Analysis (IPA) of enriched miRNAs. (A–B) IPA was performed on differentially expressed (DE) miRNAs identified in two comparisons: **A)** OPC vs non-OPC and **B)** OL vs non-OPC. **A)** DE miRNAs in OPCs relative to non-OPCs are associated with the predicted activation (red lines) of AKT, PI3K, RAS, MAP2K1/2, and insulin signaling pathways, alongside the inhibition (blue lines) of the SMAD2/3 pathway. **B)** The DE miRNAs in OL vs non-OPC comparisons are associated with predicted activation (red lines) of AKT, and inhibition (blue lines) of the estrogen receptor and SMAD2/3 pathways D-E) Predicted targets of OPC/OL-enriched miRNAs through which they potentially affect OL development and function: **C)** miR-3943, **D)** miR-488-3p, **E)** miR-675-5p, **F)** miR-4286, **G)** miR-128-3p.

In the OPC vs OL comparison, multiple miRNAs that were reduced in OPCs and enriched in OLs, including miR-22-3p, miR-135a-5p, miR-30c-5p, and miR-148a-3p, showed predicted inhibition of ERK1/2 and AKT signaling (Figure S2A). Similar to OPC vs non-OPC, downregulation of SMAD2/3 was also observed, suggesting a model in which inhibition of these pathways aid in OPC to OL maturation. Supporting this model, earlier studies have demonstrated that AKT promotes OL differentiation by phosphorylating FoxO1, which enhances expression of Sox10, a key transcription factor in OL development^39^. Similarly, the SMAD2/3 pathway, which is a downstream effector of the TGF-β signaling pathway, is known to play a complex role in OL biology. During early stages, SMAD2/3 activation can influence lineage commitment^40,41^, while at later stages it may regulate extracellular matrix remodeling, OL maturation, and interaction with astrocytes and microglia in the myelination environment^42,43^.

Next, we mapped the OPC/OL-enriched miRNAs, namely miR-3943, miR-4286, miR-1296-5p, miR-488-3p, miR-1226-3p, miR-675-5p, miR-483-5p and miR-128-3p to their predicted downstream target molecules (Figures 6C-G and 2S.B-G). This analysis revealed the direct or indirect regulation of several common targets such as ZNF488, CNSK2B, OLIG2 and SOX10 by some of these miRNAs. Notably, ASPA and TPPP were identified as targets of miR-488, KCNJ10 and FOXO1 as miR128-3p targets, LIF as a target of miR-4286, and specific inhibition of DLX1 by miR-675-5p (Figures 6 and S2). While majority of these genes have been implicated in OL biology, their roles in the context miRNA study, particularly within human cell system remain less studied, making them promising candidates as potential key molecular intermediates through which these miRNAs may influence OL development and maturation^44,45,46,47,48,49^.

## Discussion

Since the discovery of miRNA in *C.elegans* in 1993^50^ and the subsequent demonstration that they are evolutionarily conserved across species, it has become clear that they control temporal transitions during development across animal phylogenies^51^. This discovery has sparked an explosion of research focused on miRNAs, leading to the recognition of their crucial roles in development and organogenesis, as well as in the pathogenesis of numerous diseases, including cancer, cardiovascular disease, and neurodegenerative disorders^52^. An increasing number of studies have investigated the role of miRNAs in the development and function of CNS cell types, including OPCs and OLs. These studies show that several miRNAs are differentially expressed at distinct stages of OL differentiation, from progenitor cells to mature oligodendrocytes. Notably, miR-219 and miR-338 have been identified as key regulators of OL differentiation by targeting inhibitors of OL maturation and myelination pathways^8,9,53^. miR-219 is particularly important for both early and late stages of differentiation, while miR-338 functions primarily during the maturation phase, often acting synergistically with miR-219. Furthermore, overexpression of miR-219 has been reported to promote remyelination in demyelination models^8,9^. miRNA expression profiling during OL development has revealed additional miRNAs including miR-23amiR-27a, and miR-29a, which appear to fine-tune the timing of OL maturation and myelin production^54,55^. Other miRNAs such as miR-124 and miR-9 have also been implicated in maintaining the balance between OPC proliferation and differentiation by regulating transcription factors and signaling pathways essential for OL lineage progression^33,56,57^. In addition to development, changes in the miRNA profiles of neurons, microglia, OPCs, and OLs in response to disease and disease-associated stressors have also been reported^58,59^. For example, altered levels of miR-146a and miR-155 have been associated with multiple sclerosis (MS), indicating their potential roles in inflammatory responses affecting OLs^30,60^.

However, it is important to note that the majority of studies examining miRNA roles in OL lineage cell (OLLC) development have been conducted using rodent models. Given the fundamental molecular differences between human and rodent OPCs and OLs^10-12^, there is a critical need to investigate the miRNAs that regulate human OL development, lineage specification, and function. This is particularly important for improved disease modeling and for supporting therapeutic discovery in demyelinating diseases. To our knowledge, only one published study has profiled global miRNA expression in human ESC-derived OLLCs^13^. However, this study did not detect miR-219 or miR-338, two of the most consistently implicated miRNAs in OL differentiation and known to be highly enriched in primary human OPCs and OLs^9,13^, in any of their OLLC populations. Moreover, no follow-up studies have validated or further characterized the OPC/OL-associated miRNAs identified in that study. Given that the differentiation methods used at the time were less robust, it is likely that several false positives were discovered in this study. Additionally, the microarray and NanoString nCounter platforms used in many prior miRNAs profiling studies, including this study, are limited by their reliance on predefined probe sets, which restricts the ability to detect novel miRNAs or low-abundance RNA molecules^17^. In contrast, NGS-based methods offer a more comprehensive and dynamic approach to miRNA profiling, with higher sensitivity and lower false-positive rates compared to microarrays^61,62^. For this reason, we employed our human ESC reporter system^14,15^ that enables us to track and isolate well-characterized hOPCs and OLs, combined with an improved hOPC/OL differentiation protocol, to collect OLLCs at various developmental stages. We then performed miRNA profiling using an NGS-based approach (Qiagen miRNA library prep and Illumina HiSeq), which allows for a more detailed and accurate expression analysis.

Comprehensive miRNA profiling during the differentiation of hOLLCs using our reporter cell line identified several previously known as well as novel miRNAs with potential roles in OLLC differentiation. We observed the enrichment of well-characterized miRNAs at specific stages, including miR-302b-3p at the stem cell stage^20,21^; miR-26b-5p^23^ and miR-103a-3p in NPCs^24^; and miR-16-5p, miR-20a-5p, and miR-34a-5p in early OPCs^28,29,30^. miR-181a-5p, which has been implicated in both neural and oligodendrocyte development as well as in the regulation of myelination-related genes^35,36^, showed moderate expression during early differentiation, with levels gradually increasing and peaking at days 75 and 95. Importantly, the expression of stage-specific marker genes (Figure 3), together with the OPC- and OL-specific enrichment of miR-219a-5p, miR-338-3p, and miR-138 (both of which are known to cooperate with miR-219 in regulating OPC and OL differentiation^8,25-27^) further confirmed the quality of our differentiated cell populations and the reliability of our dataset. Among the lesser-studied miRNAs, we observed distinct stage-specific enrichment patterns: miR-182 in pluripotent stem cells (PSCs); miR-483-3p in neural stem cells NSCs (day 8); miR-454-3p, miR-363-3p, miR-151a-3p, and miR-10a-5p in NPCs (day 20); and miR-135a-5p, miR-27b-3p, and miR-342-3p in pre-OPCs (day 40). These miRNAs may serve as useful molecular markers for identifying human CNS cell types in future studies.

We also identified several miRNAs specifically enriched in OPCs, including miR-3943, miR-1296-5p, miR-4286, miR-3615, miR-675-5p, miR-3151-5p, miR-6726-3p, and miR-577, which may play critical roles in OLLC lineage specification and differentiation. Conversely, miR-92b-5p/3p, miR-23a-3p, miR-4443, and miR-21-3p were specifically upregulated in the non-OPC (FT) population. Interestingly, several miRNAs previously reported to be involved in hESC differentiation toward the OL lineage^13^, did not show the expected expression patterns in our dataset. For example, miR-184, previously described as OL-specific, was enriched only in the NPC population and significantly downregulated in both OPCs and OLs (Figure 3B). miR-367, reported to be expressed in ESCs, NPCs, and OPCs^20, 37^, exhibited high expression in hESCs in our dataset but was undetectable in later-stage populations. Similarly, miR-663, miR-1225-5p, and miR-638, previously reported as enriched in hESC-derived OLs^13^, were not detected in any of the populations we analyzed. Given that our OPC population, isolated based on OPC-specific PDGFRA expression, has been previously validated as closely resembling primary human OPCs^15^, the miRNAs enriched in this population are likely to be more accurate. Likewise, considering the enrichment of astrocyte markers in the non-OPC FT population^15^, the miRNAs identified in this group are likely astrocyte-specific and may serve as potential markers for astroglia identity.

Our IPA-based pathway and target analysis of differentially expressed miRNAs, with particular focus on those enriched in OPCs and OLs, suggest a coordinated regulatory network in which distinct miRNAs fine-tune AKT and SMAD2/3 activity to ensure proper timing and progression of OL differentiation. Integration of these pathways with other signaling networks, including estrogen receptor and insulin signaling, highlights a broader network of metabolic and extrinsic signals influencing OL lineage progression. Additionally, several target genes identified through the IPA analysis, such as ZNF488, CSNK2B, KCNJ1, and DLX1, have been associated with neural development and myelination. For example, ZNF488 (of Zfp488) and DLX1, are transcription factors that cooperate with Olig2 to drive neural lineage specification and OL differentiation^46,49,63^. CSNK2B, which encodes the beta subunit of casein kinase 2, is a known critical regulator of cytoskeletal dynamics required for process extension during OL maturation^44,45^. Similarly, KCNJ1 (Kir7.1), a gene encoding potassium channel, may modulate membrane potential and ion homeostasis, thereby influencing OL survival and functional maturation^47,48^. These findings support the concept that miRNA-mediated regulation extends beyond individual genes to broader signaling networks, amplifying the impact of multiple small regulatory inputs to drive more precise, stage-specific outcomes in OL development.

Collectively, our findings highlight new candidate miRNAs and molecular pathways with potential therapeutic relevance, particularly regarding enhancing remyelination in demyelinating diseases such as multiple sclerosis. Future studies will focus on experimentally validating the functional roles of these miRNAs and their targets in OL biology, both in vitro and in relevant in vivo models. Such validation could pave a way for identifying novel targets and also potentially developing miRNA-based therapeutic strategies aimed at restoring myelin integrity in demyelinating conditions.

## Material and Methods

### Human pluripotent stem cells (PSCs) and culture conditions

For this study, we used RUES1 (WiCell), NIH-approved hESC line (NIH approval number: NIHhESC-10-0012). hESCs were cultured in mTesr plus (100-0276, Stem Cell Technologies) on growth factor reduced Matrigel (354230, Corning) coated plates at 37C, 10% CO2/5% O2. Accutase (A6964, Sigma-Aldrich) was used to dissociate and passage hPSC colonies. To improve single cell survival, cells were maintained in stem cell media containing 5 mM blebbistatin (B0560, Sigma-Aldrich) for the first 24 hours after passaging. Chromosomal cells were regularly tested for mycoplasma contamination (MycoAlert, Lonza) and only cells free of contamination were utilized for OPC differentiation.

### OPC and Oligodendrocyte Differentiation Protocol

Previously generated by our lab^14,15^, hESC RPD reporter cell lines (PPM) were differentiated to OPC and OL following our well-established differentiation protocol^11^. hESCs were briefly dissociated to single cells and plated on growth factor-reduced Matrigel (354230, Corning) coated plate at 200,000 cells/well of a 6-well plate and cultured in mTesr plus (100-0276, Stem Cell Technologies) at 37C, 10% CO2/ 5% O2. Neural differentiation and spinal cord patterning was induced by dual SMAD inhibition (SB431542, 10uM and LDN193189, 250 nM) and 100 nM all-trans RA^21^ two days after passaging. Differentiating cells were maintained in neural induction media supplemented with RA (100nM) and SAG (1 mM) from day 8 to day 12. Adherent cells were lifted and cultured in low-attachment plates to promote sphere aggregation at day 12. Spheres were plated into poly-L-ornithine/laminin-coated dishes in OPC/OL differentiating media supplemented with B27 (Thermo Fisher, 12587010), N2 supplement (Thermo Fisher, 17502048), PDGF-AA (221-AA-10, R&D systems), neurotrophin-3 (GF0308, Millipore Sigma), HGF (294-HG-025 R&D systems), Minocycline (Y0001930, Sigma) and T3 (T2877, Sigma) at day 30. Cells were fed with this media 2-3 times a week until the end of the differentiation process.

### Flow Cytometry and MACS Purification of the Reporter hOPCs and hOLs

Cells were dissociated into a single cell suspension for flow cytometry analysis and Magnetic-Activated Cell Sorting (MACS) purification by incubating them in accutase (A6964, Sigma-Aldrich) for ∼45 minutes. The single cell suspension was subsequently run through a ∼70 uM cell strainer (BD Biosciences), washed and resuspended in MACS buffer (Miltenyi Biotec, Auburn, CA) for MACS cell sorting or flow analysis. An SH800S Cell Sorter (Sony Biotechnology, San Jose, CA) was used for flow analysis. The number of tdTomato+ or GFP+ cells was measured using live cells. A gate was set up using WT hES cells differentiated to day 75 or day 95. With a few minor adjustments, MACS purification was performed according to the manufacturer’s instructions. After passing through cell strainer, cells were resuspended in MACS buffer, to which, CD90.2 (THY1.2) or O4 MicroBeads were added and incubated at room temperature for 15 minutes for cell binding. The cells were passed through the LS or MS magnetic column, the columns were washed thrice with MACS buffer, and the cells that were bound to the column were obtained by pushing them through using a syringe provided. The purity of tdTomato+ population was increased by passing the collected cells through a new column without additional supplementation of MicroBeads.

### miRNA extraction and quantification

Total miRNA from three independent biological replicates of stage specific cell population were isolated using the miRNeasy Mini/Micro-Kit according to the manufacturer’s instruction (QIAGEN). The concentration of the total RNA was measured using Qubit RNA HS assay kit (Invitrogen).

### miRNA library preparation

miRNA sequencing library prepared from 10ng RNA input using QIAseq miRNA library kit (Qiagen) with minor modifications. The quantity and quality of miRNA library were determined by Qubit DNA HS assay kit (Invitrogen) and fragmented bioanalyzer (JHU Core Facility).

### miRNA profiling and data analysis

Following sample collection, RNA extraction, and preparation of miRNA libraries (Figure 2), libraries were sequenced using a full lane of illumina HiSeq (Novogene) with an estimated data output of 110Gb. After 5% for Phix control, each sample was expected to achieve 11.6M paired end read (3.5Gb data). Sequence trimming, miRNA annotation, and quantification were performed using the Qiagen Geneglobe and miRge3.0 pipeline^64,65^. After trimming the adapter sequences and filtering for the minimum read length of 16, ∼80 million trimmed reads out of total ∼300 million reads average, 40 million reads per samples were achieved. Any sample with number of reads 5 SD below the average (i.e. one of three Day 8 samples) was excluded from the downstream analysis (Table S1).

### qPCR analysis

Total miRNA from each stage-specific cell population was converted to cDNA using the High-Capacity cDNA Reverse Transcription Kit (Applied Biosystems, Cat. No. 4368814). For each qPCR reaction, 1.0 µL of synthesized cDNA was diluted with 1.3 µL of nuclease-free water to prepare a 2.3 µL template mix. Each reaction contained 0.2 µL of custom-designed primers (10 µM) and 2.5 µL of SsoAdvanced Universal SYBR Green Supermix (Bio-Rad, Cat. No. 1725270). A total of 2.7 µL of the primer–supermix mixture was added per well, bringing the final reaction volume to 5.0 µL. Reactions were performed in technical triplicates on a Bio-Rad CFX96 real-time PCR system using SYBR Green detection. Gene expression levels were calculated using the ΔΔCt method, with GAPDH and ACTB used as endogenous controls.

### IPA analysis

Ingenuity Pathway Analysis (IPA, Qiagen) was used to predict and analyze the potential downstream targets and biological networks of differentially expressed miRNAs. miRNA lists were uploaded into the IPA software, and target prediction was performed based on experimentally validated interactions and high-confidence computational predictions. Core analysis was conducted to identify canonical pathways, upstream regulators, and molecular networks associated with the miRNAs. The resulting interaction networks were visualized within IPA, and key target genes were highlighted for further functional interpretation.

### Statistical analysis

Statistical analyses were performed using GraphPad Prism (version 10.6.1). Data are presented as mean ± standard error of the mean (SEM) from three biological replicates. Comparisons between multiple groups were conducted using one-way analysis of variance (ANOVA) followed by appropriate post hoc tests. A p-value of less than 0.05 was considered statistically significant.

## Author Contributions

Methodology: M.B., S.D., X.C.; Investigation: M.B., A.D., F.C.V., Formal Analysis: A.H.P., M.K.H; Resources: X.C., S.D., D.J.Z,; Data Curation and Visualization: M.B., V.N.P.; Conceptualization: X.C.; Writing-Original Draft: M.B. Writing-Review & Editing: V.N.P., X.C.; Supervision: D.J.Z., X.C.; Funding Acquisition: M.B., X.C.

## Acknowledgement

We would like to thank the Johns Hopkins School of Medicine Genetics Resource Core Facility for their assistance with quality control of miRNA libraries. We also acknowledge the Smith’s 3^rd^ Floor Writing Accountability Group (WAG) for helping us stay committed and organized throughout the writing process. This work was supported by funding from the NIH (R00EY029011, to X.C) the Gilbert Family Foundation (to X.C), an unrestricted departmental grant to the Wilmer Eye Institute from Research to Prevent Blindness, the Myelin Repair Foundation, and fellowship fundings from the Maryland Stem Cell Research Fund and an Early Investigator Award from the DOD–MSRP (to M.B).

## Notes

### Competing Interest Statement

The authors have declared no competing interest.

## References

1 Emery, B. Regulation of oligodendrocyte differentiation and myelination. Science 330, 779–782 (2010). 10.1126/science.1190927

2 Ngo, C. & Kothary, R. MicroRNAs in oligodendrocyte development and remyelination. J Neurochem 162, 310–321 (2022). 10.1111/jnc.15618

3 Hardt, R. et al. Proteomic investigation of neural stem cell to oligodendrocyte precursor cell differentiation reveals phosphorylation-dependent Dclk1 processing. Cell Mol Life Sci 80, 260 (2023). 10.1007/s00018-023-04892-8

4 Bartel, D. P. MicroRNAs: genomics, biogenesis, mechanism, and function. Cell 116, 281–297 (2004).

5 Wu, L. & Belasco, J. G. Let me count the ways: mechanisms of gene regulation by miRNAs and siRNAs. Mol Cell 29, 1–7 (2008). 10.1016/j.molcel.2007.12.010

6 Molasy, M., Walczak, A., Szaflik, J., Szaflik, J. P. & Majsterek, I. MicroRNAs in glaucoma and neurodegenerative diseases. J Hum Genet 62, 105–112 (2017). 10.1038/jhg.2016.91

7 Doghish, A. S. et al. The role of miRNAs in multiple sclerosis pathogenesis, diagnosis, and therapeutic resistance. Pathol Res Pract 251, 154880 (2023). 10.1016/j.prp.2023.154880

8 Dugas, J. C. et al. Dicer1 and miR-219 Are required for normal oligodendrocyte differentiation and myelination. Neuron 65, 597–611 (2010). 10.1016/j.neuron.2010.01.027

9 Zhao, X. et al. MicroRNA-mediated control of oligodendrocyte differentiation. Neuron 65, 612–626 (2010). 10.1016/j.neuron.2010.02.018

10 Sim, F. J. et al. CD140a identifies a population of highly myelinogenic, migration-competent and efficiently engrafting human oligodendrocyte progenitor cells. Nature biotechnology 29, 934–941 (2011). 10.1038/nbt.1972

11 Sim, F. J., Windrem, M. S. & Goldman, S. A. Fate determination of adult human glial progenitor cells. Neuron Glia Biol 5, 45–55 (2009). 10.1017/S1740925X09990317

12 Sim, F. J. et al. Complementary patterns of gene expression by human oligodendrocyte progenitors and their environment predict determinants of progenitor maintenance and differentiation. Annals of neurology 59, 763–779 (2006). 10.1002/ana.20812

13 Letzen, B. S. et al. MicroRNA expression profiling of oligodendrocyte differentiation from human embryonic stem cells. PLoS One 5, e10480 (2010). 10.1371/journal.pone.0010480

14 Li, W. et al. High-Throughput Screening for Myelination Promoting Compounds Using Human Stem Cell-derived Oligodendrocyte Progenitor Cells Identifies Novel Targets. bioRxiv, 2022.2001.2018.476755 (2022). 10.1101/2022.01.18.476755

15 Chamling, X. et al. Single-cell transcriptomic reveals molecular diversity and developmental heterogeneity of human stem cell-derived oligodendrocyte lineage cells. Nature communications 12, 652 (2021). 10.1038/s41467-021-20892-3

16 Douvaras, P. & Fossati, V. Generation and isolation of oligodendrocyte progenitor cells from human pluripotent stem cells. Nat Protoc 10, 1143–1154 (2015). 10.1038/nprot.2015.075

17 Godoy, P. M. et al. Comparison of Reproducibility, Accuracy, Sensitivity, and Specificity of miRNA Quantification Platforms. Cell Rep 29, 4212–4222.e4215 (2019). 10.1016/j.celrep.2019.11.078

18 Liu, J., Jennings, S. F., Tong, W. & Hong, H. Next generation sequencing for profiling expression of miRNAs: technical progress and applications in drug development. J Biomed Sci Eng 4, 666–676 (2011). 10.4236/jbise.2011.410083

19 Buermans, H. P., Ariyurek, Y., van Ommen, G., den Dunnen, J. T. & t Hoen, P. A. New methods for next generation sequencing based microRNA expression profiling. BMC Genomics 11, 716 (2010). 10.1186/1471-2164-11-716

20 Yang, S. L. et al. MiR-302/367 regulate neural progenitor proliferation, differentiation timing, and survival in neurulation. Dev Biol 408, 140–150 (2015). 10.1016/j.ydbio.2015.09.020

21 Parchem, R. J. et al. miR-302 Is Required for Timing of Neural Differentiation, Neural Tube Closure, and Embryonic Viability. Cell Rep 12, 760–773 (2015). 10.1016/j.celrep.2015.06.074

22 Subramanyam, D. et al. Multiple targets of miR-302 and miR-372 promote reprogramming of human fibroblasts to induced pluripotent stem cells. Nat Biotechnol 29, 443–448 (2011). 10.1038/nbt.1862

23 Sauer, M. et al. The miR-26 family regulates neural differentiation-associated microRNAs and mRNAs by directly targeting REST. J Cell Sci 134 (2021). 10.1242/jcs.257535

24 Annibali, D. et al. A new module in neural differentiation control: two microRNAs upregulated by retinoic acid, miR-9 and -103, target the differentiation inhibitor ID2. PLoS One 7, e40269 (2012). 10.1371/journal.pone.0040269

25 Nazari, B. et al. Overexpression of miR-219 promotes differentiation of human induced pluripotent stem cells into pre-oligodendrocyte. J Chem Neuroanat 91, 8–16 (2018). 10.1016/j.jchemneu.2018.03.001

26 Fan, H. B. et al. Transplanted miR-219-overexpressing oligodendrocyte precursor cells promoted remyelination and improved functional recovery in a chronic demyelinated model. Sci Rep 7, 41407 (2017). 10.1038/srep41407

27 Wang, H. et al. miR-219 Cooperates with miR-338 in Myelination and Promotes Myelin Repair in the CNS. Developmental cell 40, 566–582 e565 (2017). 10.1016/j.devcel.2017.03.001

28 Arzhanov, I., Sintakova, K. & Romanyuk, N. The Role of miR-20 in Health and Disease of the Central Nervous System. Cells 11 (2022). 10.3390/cells11091525

29 Hermeking, H. The miR-34 family in cancer and apoptosis. Cell Death Differ 17, 193–199 (2010). 10.1038/cdd.2009.56

30 Junker, A. et al. MicroRNA profiling of multiple sclerosis lesions identifies modulators of the regulatory protein CD47. Brain 132, 3342–3352 (2009). 10.1093/brain/awp300

31 Zhao, C., Sun, G., Li, S. & Shi, Y. A feedback regulatory loop involving microRNA-9 and nuclear receptor TLX in neural stem cell fate determination. Nat Struct Mol Biol 16, 365–371 (2009). 10.1038/nsmb.1576

32 Buller, B. et al. Regulation of serum response factor by miRNA-200 and miRNA-9 modulates oligodendrocyte progenitor cell differentiation. Glia 60, 1906–1914 (2012). 10.1002/glia.22406

33 Lau, P. et al. Identification of dynamically regulated microRNA and mRNA networks in developing oligodendrocytes. The Journal of neuroscience : the official journal of the Society for Neuroscience 28, 11720–11730 (2008). 10.1523/JNEUROSCI.1932-08.2008

34 Lecca, D. et al. MiR-125a-3p timely inhibits oligodendroglial maturation and is pathologically up-regulated in human multiple sclerosis. Sci Rep 6, 34503 (2016). 10.1038/srep34503

35 Maciak, K., Dziedzic, A. & Saluk, J. Remyelination in multiple sclerosis from the miRNA perspective. Front Mol Neurosci 16, 1199313 (2023). 10.3389/fnmol.2023.1199313

36 Pietrasik, S., Dziedzic, A., Miller, E., Starosta, M. & Saluk-Bijak, J. Circulating miRNAs as Potential Biomarkers Distinguishing Relapsing-Remitting from Secondary Progressive Multiple Sclerosis. A Review. Int J Mol Sci 22 (2021). 10.3390/ijms222111887

37 Ghasemi-Kasman, M., Zare, L., Baharvand, H. & Javan, M. In vivo conversion of astrocytes to myelinating cells by miR-302/367 and valproate to enhance myelin repair. J Tissue Eng Regen Med 12, e462–e472 (2018). 10.1002/term.2276

38 Hou, H., Wang, Y., Yang, L. & Wang, Y. Exosomal miR-128-3p reversed fibrinogen-mediated inhibition of oligodendrocyte progenitor cell differentiation and remyelination after cerebral ischemia. CNS Neurosci Ther 29, 1405–1422 (2023). 10.1111/cns.14113

39 Wang, H. et al. Akt Regulates Sox10 Expression to Control Oligodendrocyte Differentiation via Phosphorylating FoxO1. J Neurosci 41, 8163–8180 (2021). 10.1523/jneurosci.2432-20.2021

40 Palazuelos, J., Klingener, M. & Aguirre, A. TGFβ signaling regulates the timing of CNS myelination by modulating oligodendrocyte progenitor cell cycle exit through SMAD3/4/FoxO1/Sp1. J Neurosci 34, 7917–7930 (2014). 10.1523/jneurosci.0363-14.2014

41 Hamaguchi, M. et al. Circulating transforming growth factor-β1 facilitates remyelination in the adult central nervous system. Elife 8 (2019). 10.7554/eLife.41869

42 Baror, R. et al. Transforming growth factor-beta renders ageing microglia inhibitory to oligodendrocyte generation by CNS progenitors. Glia 67, 1374–1384 (2019). 10.1002/glia.23612

43 Doyle, K. P., Cekanaviciute, E., Mamer, L. E. & Buckwalter, M. S. TGFβ signaling in the brain increases with aging and signals to astrocytes and innate immune cells in the weeks after stroke. J Neuroinflammation 7, 62 (2010). 10.1186/1742-2094-7-62

44 Huillard, E. et al. Disruption of CK2beta in embryonic neural stem cells compromises proliferation and oligodendrogenesis in the mouse telencephalon. Mol Cell Biol 30, 2737–2749 (2010). 10.1128/mcb.01566-09

45 Yang, C. P. et al. Comprehensive integrative analyses identify GLT8D1 and CSNK2B as schizophrenia risk genes. Nat Commun 9, 838 (2018). 10.1038/s41467-018-03247-3

46 Soundarapandian, M. M. et al. Zfp488 promotes oligodendrocyte differentiation of neural progenitor cells in adult mice after demyelination. Sci Rep 1, 2 (2011). 10.1038/srep00002

47 Papanikolaou, M., Butt, A. M. & Lewis, A. A critical role for the inward rectifying potassium channel Kir7.1 in oligodendrocytes of the mouse optic nerve. Brain Struct Funct 225, 925–934 (2020). 10.1007/s00429-020-02043-4

48 Larson, V. A. et al. Oligodendrocytes control potassium accumulation in white matter and seizure susceptibility. Elife 7 (2018). 10.7554/eLife.34829

49 Petryniak, M. A., Potter, G. B., Rowitch, D. H. & Rubenstein, J. L. Dlx1 and Dlx2 control neuronal versus oligodendroglial cell fate acquisition in the developing forebrain. Neuron 55, 417–433 (2007). 10.1016/j.neuron.2007.06.036

50 Lee, R. C., Feinbaum, R. L. & Ambros, V. The C. elegans heterochronic gene lin-4 encodes small RNAs with antisense complementarity to lin-14. Cell 75, 843–854 (1993). 10.1016/0092-8674(93)90529-y

51 Pasquinelli, A. E. et al. Conservation of the sequence and temporal expression of let-7 heterochronic regulatory RNA. Nature 408, 86–89 (2000). 10.1038/35040556

52 Bhaskaran, M. & Mohan, M. MicroRNAs: history, biogenesis, and their evolving role in animal development and disease. Vet Pathol 51, 759–774 (2014). 10.1177/0300985813502820

53 Wang, H. et al. miR-219 Cooperates with miR-338 in Myelination and Promotes Myelin Repair in the CNS. Dev Cell 40, 566–582.e565 (2017). 10.1016/j.devcel.2017.03.001

54 Lin, S. T. & Fu, Y. H. miR-23 regulation of lamin B1 is crucial for oligodendrocyte development and myelination. Dis Model Mech 2, 178–188 (2009). 10.1242/dmm.001065

55 Tripathi, A. et al. Oligodendrocyte Intrinsic miR-27a Controls Myelination and Remyelination. Cell Rep 29, 904–919.e909 (2019). 10.1016/j.celrep.2019.09.020

56 Zhang, W. H., Jiang, L., Li, M. & Liu, J. MicroRNA-124: an emerging therapeutic target in central nervous system disorders. Exp Brain Res 241, 1215–1226 (2023). 10.1007/s00221-022-06524-2

57 Ma, Q., Zhang, L. & Pearce, W. J. MicroRNAs in brain development and cerebrovascular pathophysiology. Am J Physiol Cell Physiol 317, C3–c19 (2019). 10.1152/ajpcell.00022.2019

58 Nguyen, T. P. N., Kumar, M., Fedele, E., Bonanno, G. & Bonifacino, T. MicroRNA Alteration, Application as Biomarkers, and Therapeutic Approaches in Neurodegenerative Diseases. Int J Mol Sci 23 (2022). 10.3390/ijms23094718

59 Ma, Q., Matsunaga, A., Ho, B., Oksenberg, J. R. & Didonna, A. Oligodendrocyte-specific Argonaute profiling identifies microRNAs associated with experimental autoimmune encephalomyelitis. J Neuroinflammation 17, 297 (2020). 10.1186/s12974-020-01964-5

60 Moore, C. S. et al. miR-155 as a multiple sclerosis-relevant regulator of myeloid cell polarization. Ann Neurol 74, 709–720 (2013). 10.1002/ana.23967

61 Motameny, S., Wolters, S., Nürnberg, P. & Schumacher, B. Next Generation Sequencing of miRNAs - Strategies, Resources and Methods. Genes (Basel) 1, 70–84 (2010). 10.3390/genes1010070

62 Landgraf, P. et al. A mammalian microRNA expression atlas based on small RNA library sequencing. Cell 129, 1401–1414 (2007). 10.1016/j.cell.2007.04.040

63 Leung, R. F. et al. Genetic Regulation of Vertebrate Forebrain Development by Homeobox Genes. Frontiers in Neuroscience Volume 16 - 2022 (2022). 10.3389/fnins.2022.843794

64 Patil, A. H. & Halushka, M. K. miRge3.0: a comprehensive microRNA and tRF sequencing analysis pipeline. NAR Genom Bioinform 3, lqab068 (2021). 10.1093/nargab/lqab068

65 Patil, A. H., Baran, A., Brehm, Z. P., McCall, M. N. & Halushka, M. K. A curated human cellular microRNAome based on 196 primary cell types. Gigascience 11 (2022). 10.1093/gigascience/giac083

