## Supplementary figures and tables for "Profiling miRNAs involved in Human Oligodendrocyte Precursor Cell Differentiation and Maturation"

**Supplementary Materials**

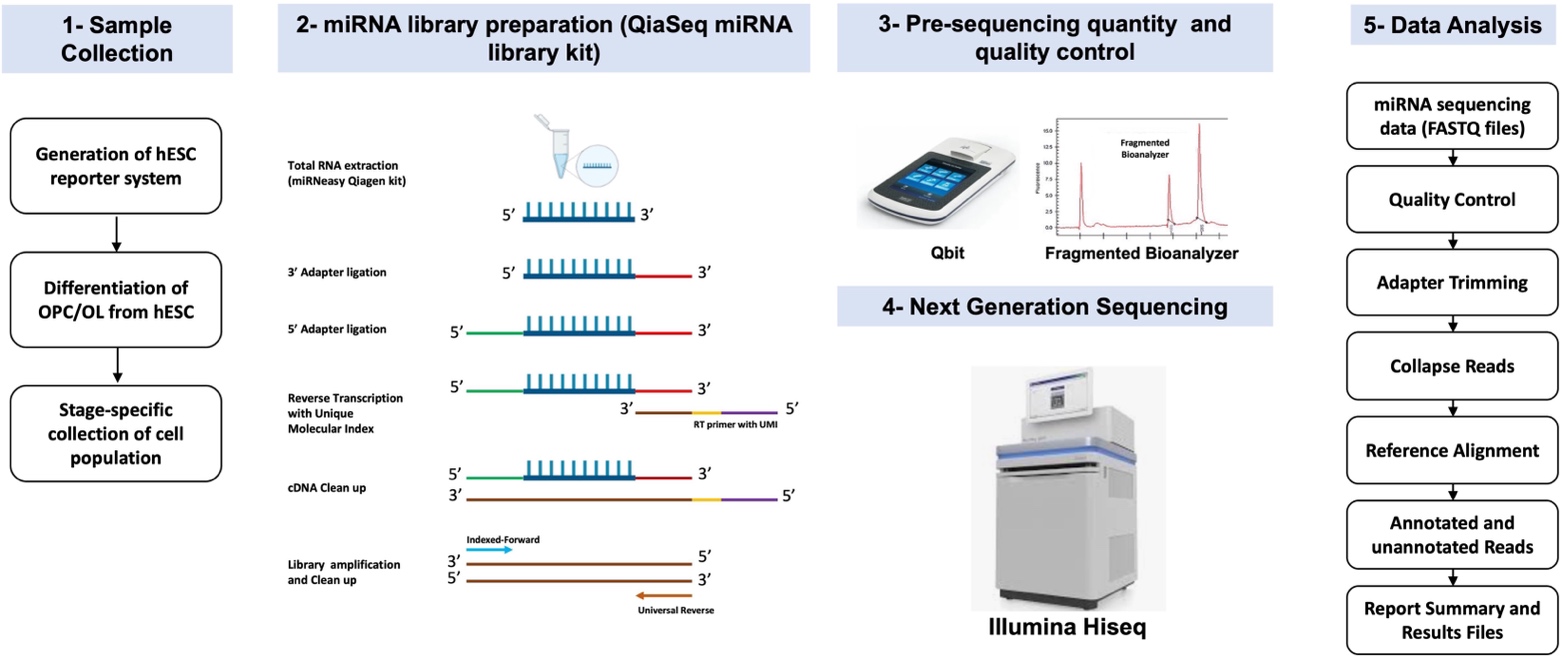

**Figure S1. Schematic overview of the approaches. 1.** Sample preparation and collection; **2.** miRNA library preparation; **3.** Quantity and quality control of the miRNA libraries using Q-bit and fragmented bioanalyzer, respectively; **4.** Next generation sequencing; and **5.** Data analysis pipeline.

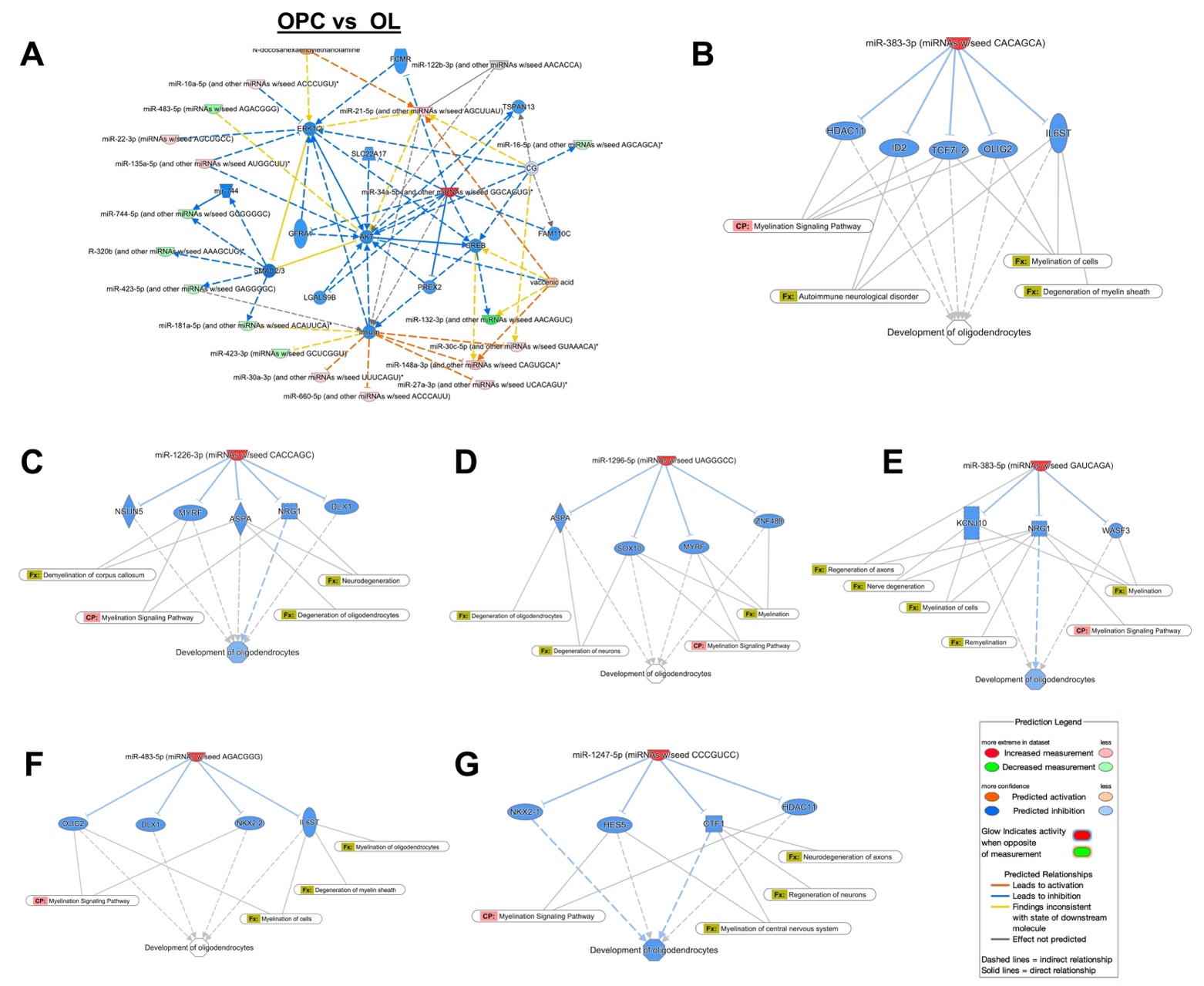

**Figure S2. Ingenuity Pathway Analysis (IPA) of enriched miRNAs**. **A)** IPA was performed on differentially expressed (DE) miRNAs identified in OL vs OPC comparison, which demonstrated predicted inhibition of ERK1/2, AKT, and SMAD2/3 signaling. B-G) Predicted mapped targets of OPC/OL-enriched miRNAs through which they potentially affect OL development and function: **B)** miR-383-5p, **C)** miR-1226-3p, **D)** miR-1296, **E)** miR-383-5p, **F)** miR-483-5p, and **G)** miR-1247-5p.

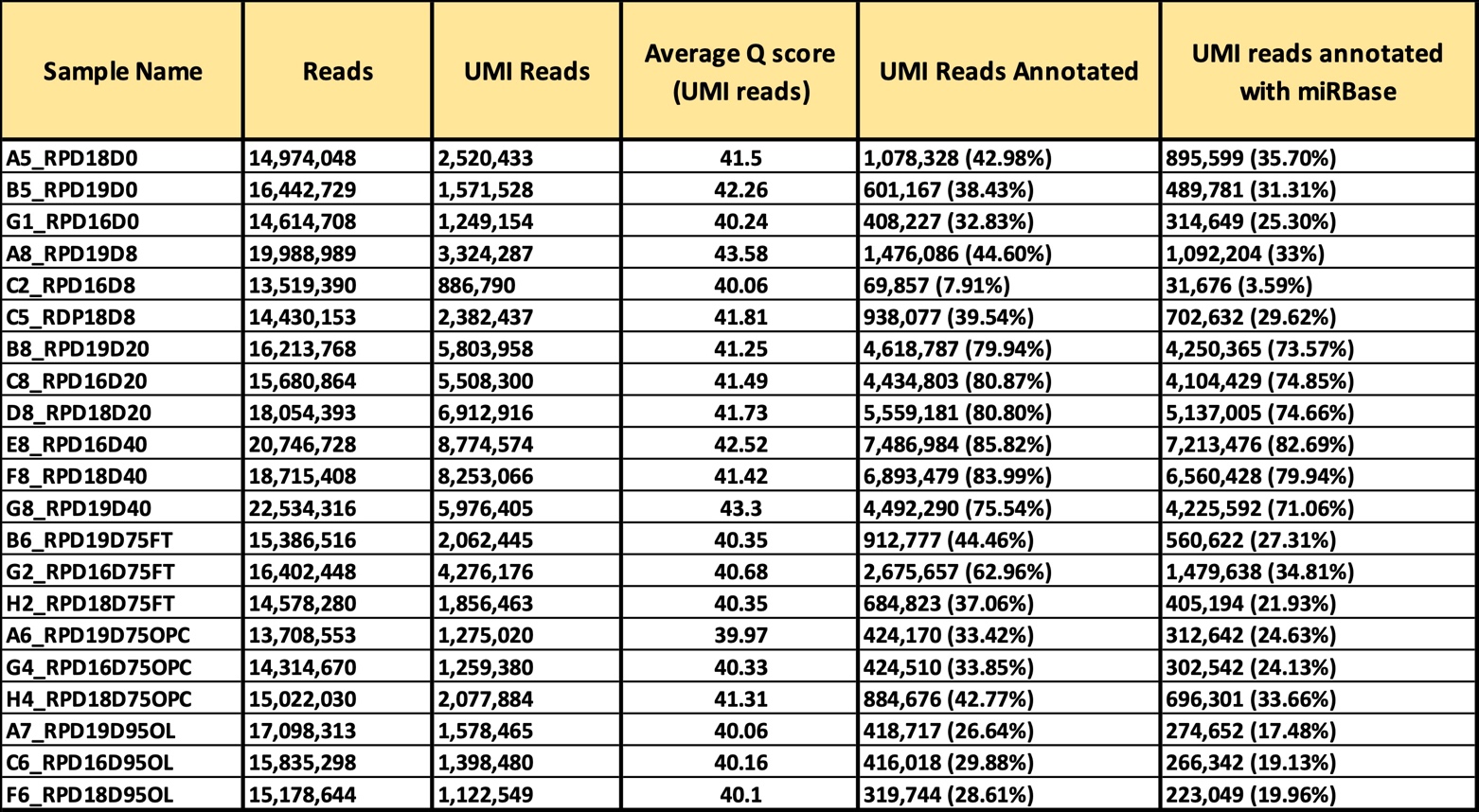

**Table S1. Quality control metrics for miRNA sequencing libraries.** A table summarizing sequencing and quality control metrics for each miRNA library, including total reads, UMI-filtered reads, average Q scores for UMI reads, the number of UMI reads successfully annotated, and the subset of annotated UMI reads mapped to miRBase. These metrics were generated by Qiagen global analysis portal.

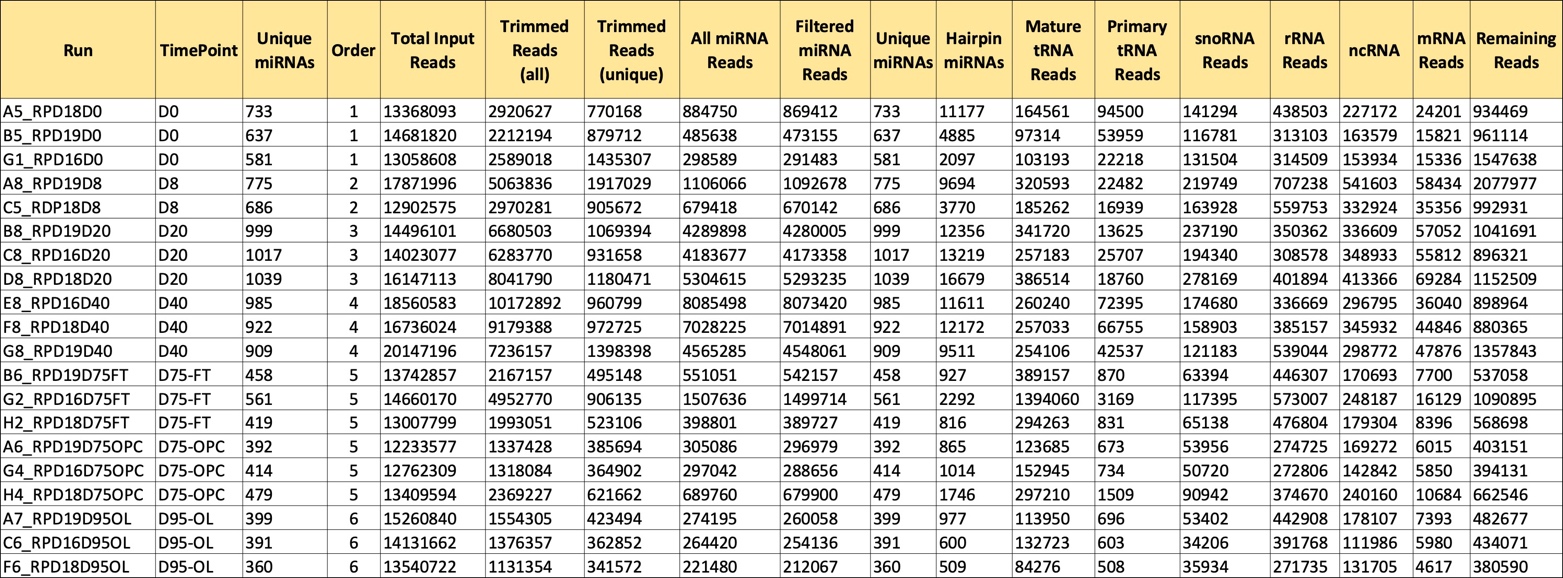

**Table S2. Comprehensive annotation and processing metrics for miRNA sequencing data using miRge3.0 analysis tool.** A table summarizing read processing and annotation outputs for each sample, including total input reads, trimmed reads (all and unique), total miRNA reads, and filtered miRNA reads. It also reports the distribution of annotated RNA species, including unique miRNAs, hairpin miRNAs, mature tRNA reads, primary tRNA reads, snoRNA reads, rRNA reads, other non-coding RNAs, mRNA reads, and remaining unclassified reads. These metrics provide an overview of library composition and sequencing quality.

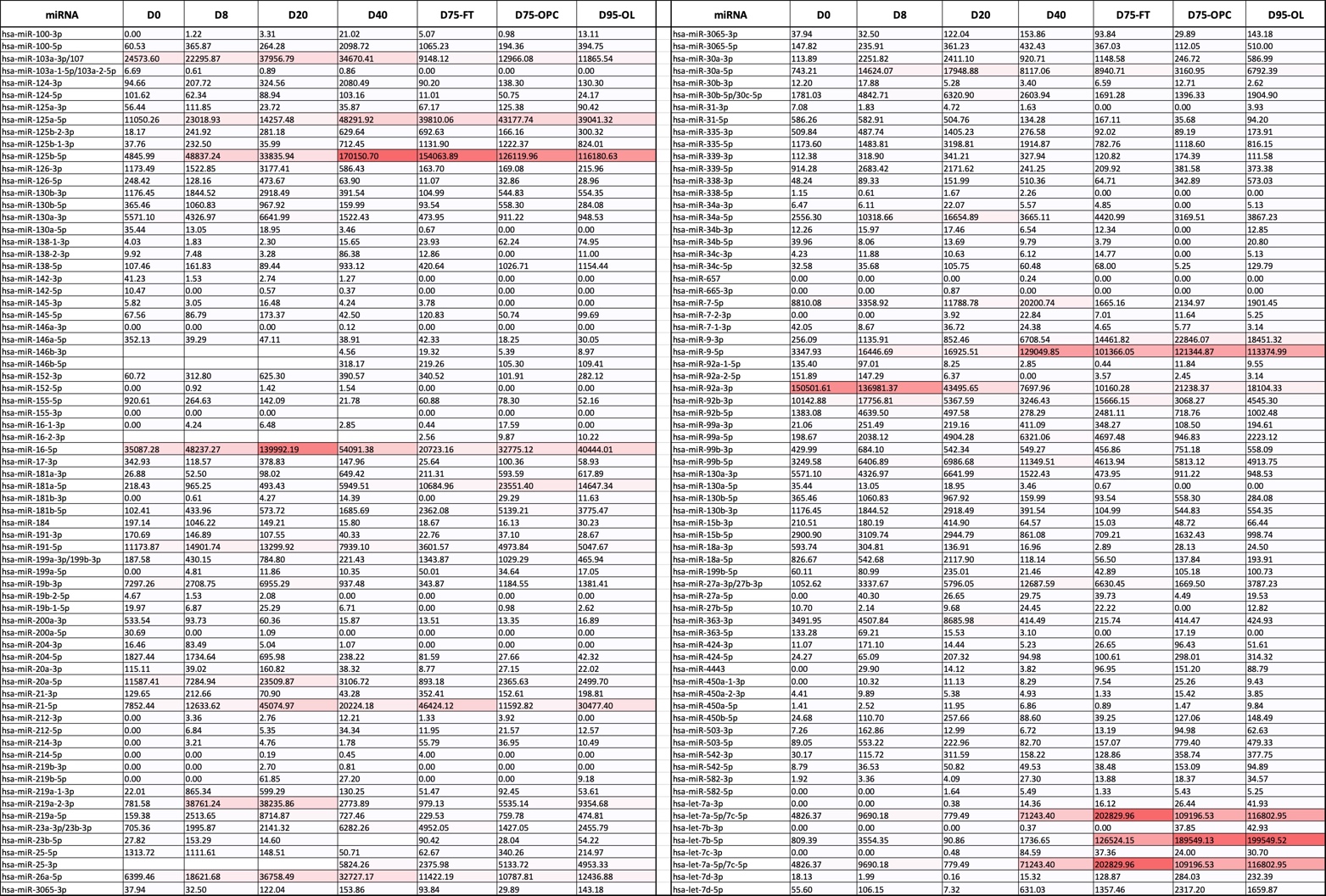

**Table S3.** Average of RPM values for previously known miRNAs at different stages of OL differentiation.
